# Identification of High risk nsSNPs in Human *TP53* Gene Associated with Li–Fraumeni Syndrome: An *In Silico* Analysis Approach

**DOI:** 10.1101/2020.12.04.411835

**Authors:** Mujahed I. Mustafa, Naseem S. Murshed, Mazen A. Elbasher, Abdelrafie M. Makhawi

**Affiliations:** Department of Biotechnology, University of Bahri, Khartoum, Sudan; Department of Microbiology, International University of Africa, Khartoum, Sudan; Department of Chemistry, University of Bahri, Khartoum, Sudan

**Author notes:** Correspondence should be addressed to Mujahed I. Mustafa.

**Keywords:** *In Silico* analysis, *TP53*, Diagnostic markers, Li–Fraumeni syndrome, single nucleotide polymorphisms

## Abstract

**Background:** Li–Fraumeni syndrome (LFS) is a cancer–prone conditions caused by a germline mutation of the TP53 gene on chromosome 17p13.1. It has an autosomal dominant pattern of inheritance with high penetrance.

**Purpose:** The aim of this study is to identify the high-risk pathogenic nsSNPs in *PT53* gene that could be involved in the pathogenesis of Li–Fraumeni syndrome.

**Methods:** The nsSNPs in the human *PT53* gene retrieved from NCBI, were analyzed for their functional and structural consequences using various *in silico* tools to predict the pathogenicity of each SNP. SIFT, Polyphen, PROVEAN, SNAP2, SNPs&Go, PHD-SNP, and P-Mut were chosen to study the functional inference while I-Mutant 3.0, and MUPro tools were used to test the impact of amino acid substitutions on protein stability by calculating ΔΔG value. The effects of the mutations on 3D structure of the *PT53* protein were predicted using RaptorX and visualized by UCSF Chimera.

**Results:** A total of 845 *PT53* nsSNPs were analyzed. Out of 7 nsSNPs of *PT53* three of them (T118L, C242S, and I251N) were found high-risk pathogenic.

**Conclusion:** In this study, out of 7 predicted high-risk pathogenic nsSNPs, three high-risk pathogenic nsSNPs of *PT53* gene were identified, which could be used as diagnostic marker for this gene. The combination of sequence-based and structure-based approaches is highly effective for pointing pathogenic regions.

## 1 Introduction

Li–Fraumeni syndrome (LFS) is a cancer–prone conditions caused by a germline mutation of the TP53 (tumor protein 53) gene on chromosome 17p13.1 [1, 2]. first described in 1969 by Li and Fraumeni [3]. It has an autosomal dominant pattern of inheritance with high penetrance. LFS related typically with sarcomas, premenopausal breast cancer, brain tumors, and adrenocortical carcinomas [3-6], which it makes it one of the most devastating cancer predisposition syndromes [7]. LFS has long been seen as a rare syndrome but recent studies proves otherwise. The prevalence of LFS is about 1:5000 with a high interregional variance [7].

Two published algorithms are utilized to identify patients at risk of LFS who would benefit from molecular testing, the classical LFS criteria and the Chompret criteria [8, 9]. A *TP53* germline mutation was found in 29%and 35% of French and American families, respectively, who fulfilled the 2001 Chompret criteria [10, 11].

80% of the LFS cases had been reported due to germline *TP53* mutations. since approximately 20 % of LFS families do not exhibit TP53 mutations [4, 12, 13].The underlying genetic mutations in these families remain to be discovered. Yet, de novo mutations occur in ∼10%–20% of LFS cases [3, 14].

Recent studies have identified large gene deletions in some families negative for point mutations [15, 16] Despite intensive search, no other gene has been significantly associated with LFS. Reports that germline mutations in the *CHEK2* gene may predispose to LFS have not been substantiated [17, 18]. Germline mutations in *CHEK2*, including the common 1100delC frameshift mutation in exon 10, are currently considered as low-penetrance mutations associated with a three to four fold increase in the risk of breast cancer [19], which is one of the most frequent cancer in adult patients with LFS [20-22].

The PT53 protein is a transcription factor expressed in most cell types, upregulating the transcription of target genes involved in cell cycle arrest, DNA repair, apoptosis, and senescence, in response to DNA damage [23]. The vital role of TP53 mutation in causing LFS is highlighted by the point that mice lacking functional trp53 alleles develop lymphomas and sarcomas. Knock-in mice with germline missense mutations in trp53 simulating LFS develop more destructive tumors as matched with mice lacking trp53 alleles, suggesting that dominant gain of-function effects [24].

Disease-causing single nucleotide polymorphisms (SNPs) are frequently found to arise at evolutionarily conserved regions; these have a key role at structural and functional levels of the protein [25, 26]. The capability to calculate whether a particular SNP is deleterious or not is very important for the prognosis of disorder. SNPs may or may not affect protein function [27]. Therefore, it is necessary to understand the association between the SNPs and its phenotypic influence which might be useful in analyzing the reasons of numerous diseases or disorders. The single most frequent mutation is p.R337H, an atypical variant of incomplete penetrance which is present at high prevalence in the population of Southern Brazil due to a widespread founder effect [28-30].

Lately, screening of SNPs by the computational approach is very hot topic in revealing the mutations that are affects protein structure and function [31]. The nsSNPs of the *PT53* gene has not been considered to date to observe their functional and structural effects on the molecular level. To address this matter, this study therefore used several bioinformatics tools for the identification of pathogenic SNPs in the coding region of *PT53* gene, which could be used as diagnostic markers for this devastating syndrome.

## 2 Materials and Methods

### 2.1 nsSNPs Data retrieval

The nsSNPs data were retrieved from SNP database (http://www.ncbi.nlm.nih.gov/projects/SNP/) while the protein reference sequence (ID: P04637) was retrieved from Uniprot database [32].

### 2.2 Prediction of nsSNPs Pathogenicity

#### 2.2.1. SIFT

It is the first in silico functional analysis that calculates whether an amino acid alteration affects protein function or not. Sorting Intolerant From Tolerant (SIFT) scores < 0.05 are expected to be damaging altered amino acid, otherwise it is considered to be tolerant [33].

#### 2.2.2. Polyphen

It is a trained machine learning to predict whether an amino acid replacement affects protein function and structure or not, by calculating position-specific independent count (PSIC) for each SNP at a time. There are 2 outputs whether probably damaging (values are more frequently 1) and possibly damaging or benign (values range from 0 to 0.95) [34].

#### 2.2.3. PROVEAN

It is an online in silico functional analysis tool that calculates whether an amino acid replacement has an influence on the organic function of a protein stranded on the alignment-based score. If the PROVEAN score ⩽–2.5, the protein variant is expected to have a “deleterious” effect, whereas if the PROVEAN score is >–2.5, the variant is expected to have a “neutral” effect [35].

#### 2.2.4. SNAP2

It is a trained functional analysis tool that differentiates between effect and neutral SNPs by taking various features into validation. SNAP2 got an accuracy of 83%, which has 2 expectations: effect (positive score) or neutral (negative score). It is considered an important and substantial enhancement over other methods [36].

#### 2.2.5. SNPs&Go

It is a trained machine learning based on the technique to precisely calculates the deleterious associated alterations from protein sequence. SNPs&Go collects in a unique framework information derived from protein sequence, evolutionary information, and function as coded in the Gene Ontology terms and underperforms other available predictive method (PhD-SNP) [37].

#### 2.2.6. P-Mut

It is a web-based tool for the explanation of SNP alternates on proteins, which allows the rapid and precise calculation (80%) of the compulsive features of each SNP grounded on the practice of neural networks [38].

### 2.3. Protein Stability Prediction

#### 2.3.1. I-Mutant 3.0

I-Mutant is a support vector machine (SVM)-based tool. I-Mutant predicts whether the protein mutation stabilizes or destabilizes the protein structure by calculating free energy change by coupling predictions with the energy-based FOLD-X tool [39].

#### 2.3.2. MUPro

It is a structural analysis online tool for the calculation of protein stability variations in arbitrary SNPs. The value of the energy change is expected, and assurance mark between –1 and 1 for evaluating the assurance of the expectation is calculated. A score of <0 means the mutant decreases the protein stability; conversely, a score of >0 means the mutant increases the protein stability [40].

### 2.4. Evolutionary conservation analysis of nsSNPs

ConSurf server It is a web server that offers evolutionary conservation summaries for proteins of known structure in the protein data bank. ConSurf spots the parallel amino acid sequences and runs multi-alignment methods. The conserved amino acid across species flags its position using specific algorithm [41].

### 2.5. Protein secondary structure prediction and interactions

PSIPRED was also used to predict and validates the secondary structure of *PT53* protein. The PSIPRED server developed by PSI-BLAST based on specific matrices. The input was the protein sequence in FASTA format, while the output was Protein secondary structures [42].

### 2.6. 3D structure prediction, visualization and physiochemical changes of substitutions

#### 2.6.1. RaptorX

It have been used to generate a 3D structural model for wildtype *PT53*. The FASTA format sequence of *PT53* protein was retrieved from UniProt; it was then used as an input to predict the 3D structure of human *PT53* protein [43].

#### 2.6.2. UCSF Chimera

It is a visualization tool of 3D structure prototype. A predicted model was formed by RaptorX to visualize and compare the wild and mutant amino acids using UCSF Chimera [44].

#### 2.6.3. HOPE

It is a server to search protein 3D structures by data mining from several sources databases. It was used to analyze the physiochemical changes induced by the mutations and to confirm the outcomes that we achieved earlier. The inputs were the protein sequence and a single mutation, while the outputs were the physiochemical changes triggered by the mutations [45].

## 3. Results

### 3.1. Prediction of deleterious SNPs of *TP53* gene

The total number of SNPs in the coding region of *TP53* gene that were retrieved from dbSNP database were 845 nsSNPs. These SNPs were submitted into diverse functional analysis tools **(Figure 1)**. 127 out of 845 nsSNPs were found to be affect by SIFT, 114 damaging SNPs (5 possibly damaging and 120 probably damaging) by using Polyphen 189 were found to be deleterious by PROVEAN and 157 were predicted deleterious by SNAP2 **(Table 1)**. The quad-positive deleterious SNPs from the previous 4 analysis tools, out of 845 SNPs there were 7 predicted pathogenic SNPs, and then were submitted into SNPs&Go, PHD-SNP, and P-Mut to get more accurate prediction that could effects on the functional influence; the triple positive results from the three tools were found to be 4 SNPs **(Table 2)**.

**Table 1:** Pathogenic SNPs in *TP53* gene as predicted by four computational tools:

| SUB | SIFT prediction | Score | Polyphen2 prediction | Score | PROVEAN prediction | Score | SNAP2 Prediction | Score |
| --- | --- | --- | --- | --- | --- | --- | --- | --- |
| K120P | Deleterious | 0 | Probably damaging | 1 | Deleterious | -5.644 | effect | 75 |
| I251N | Deleterious | 0 | Probably damaging | 1 | Deleterious | -6.591 | effect | 13 |
| T118L | Deleterious | 0 | Probably damaging | 1 | Deleterious | -5.028 | effect | 40 |
| G117M | Deleterious | 0 | Probably damaging | 1 | Deleterious | -4.834 | effect | 34 |
| F113V | Deleterious | 0 | Probably damaging | 1 | Deleterious | -6.155 | effect | 48 |
| F113R | Deleterious | 0 | Probably damaging | 1 | Deleterious | -7.975 | effect | 75 |
| C242S | Deleterious | 0 | Probably damaging | 1 | Deleterious | -9.759 | effect | 85 |

**Table 2:** The most damaging SNPs in *TP53* gene as predicted by three different online tools:

| SUB | SNPs&Go Prediction | RI | Probability | PHD-SNP Prediction | RI | Probability | P-Mut prediction | Probability |
| --- | --- | --- | --- | --- | --- | --- | --- | --- |
| K120P | Disease | 5 | 0.763 | Disease | 7 | 0.839 | Disease | 0.86 (91%) |
| I251N | Disease | 7 | 0.835 | Disease | 7 | 0.863 | Disease | 0.86 (91%) |
| T118L | Disease | 5 | 0.741 | Disease | 5 | 0.744 | Disease | 0.78 (88%) |
| G117M | Neutral | 1 | 0.448 | Neutral | 2 | 0.405 | Disease | 0.56 (81%) |
| F113V | Neutral | 0 | 0.484 | Disease | 7 | 0.854 | Disease | 0.78 (88%) |
| F113R | Neutral | 2 | 0.379 | Disease | 7 | 0.854 | Disease | 0.86 (91%) |
| C242S | Disease | 8 | 0.893 | Disease | 6 | 0.802 | Disease | 0.91 (93%) |

**Table 3:** Shows protein stabilities of *TP53* nsSNPs by I-Mutant 3.0 and MUPro:

| SUB | I-Mutant Prediction | RI | DDG Value Prediction | MUPro Prediction | MUPro Score |
| --- | --- | --- | --- | --- | --- |
| K120P | Increase | 6 | 0.01 | Decrease | -0.67801266 |
| I251N | Decrease | 5 | -1.99 | Decrease | -1.9395329 |
| T118L | Decrease | 6 | -0.73 | Decrease | -0.54246124 |
| C242S | Decrease | 7 | -1.13 | Decrease | -2.8407208 |

**Figure 1:**
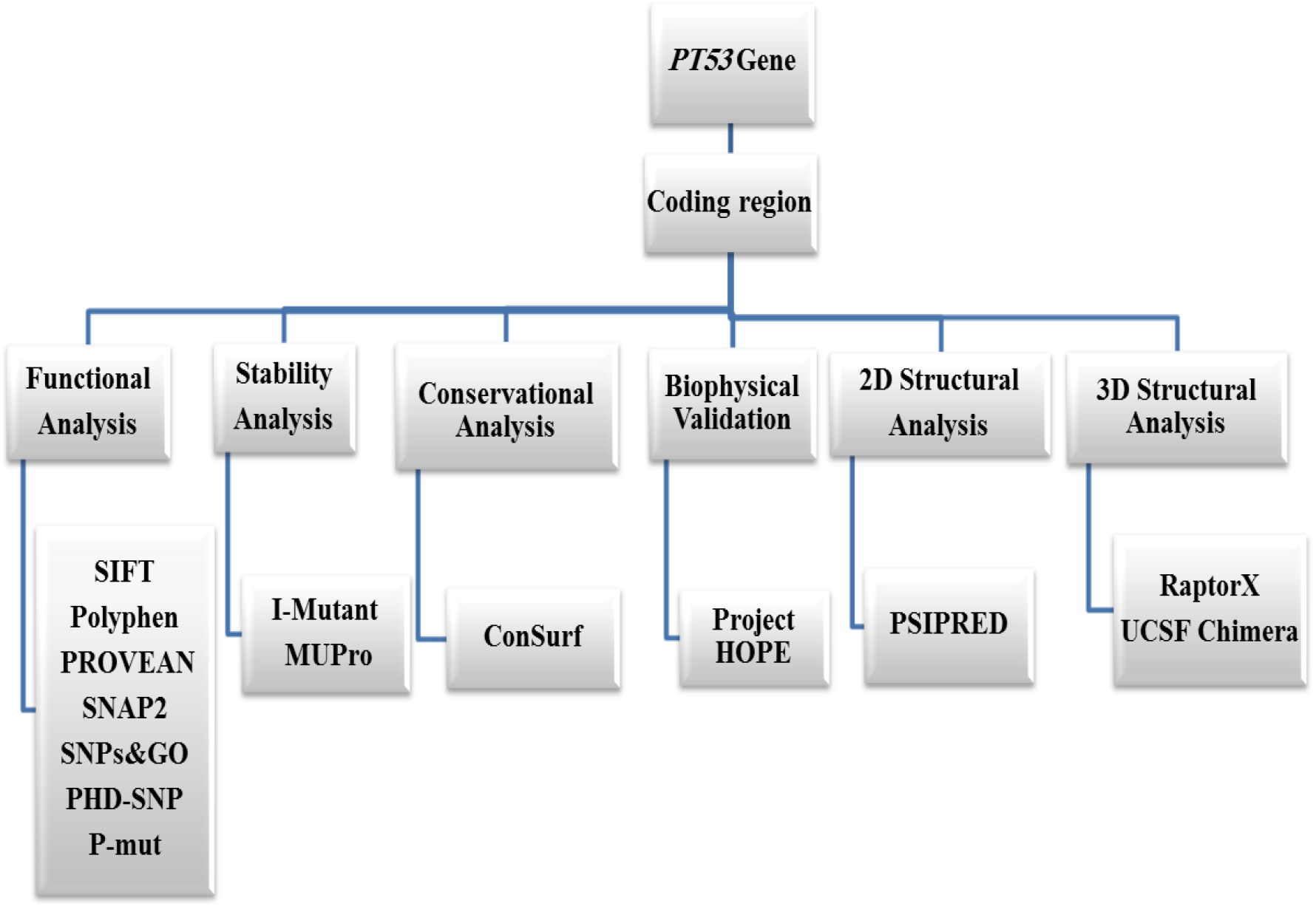
Graphic representation of *TP53* gene work flow.

### 3.2. Prediction of protein structural stability

The protein stability analysis with these 4 SNPs (T118L, K120, C242S, and I251N) was performed by using I-Mutant and MUPro, which discovered that all SNPs decreased the protein stability except K120P, which increased the protein stability. All further analysis was hold for these SNPs (T118L, C242S, and I251N).

### 3.3. Evolutionary conservation analysis of nsSNPs

The conservation regions of TP53 protein had been predicted by Consurf server; the result shows 3 SNPs (T118L, C242S, and I251N) located in highly conserved regions.

### 3.4. Protein secondary structure prediction and interactions

The secondary structure of TP53 was evaluated by PSIPRED server. The results revealed a mixture of coil, alpha helix and extended strand. The random coil was observed to be the main secondary structural motif (63.61%), followed by Alpha helix (19.34%) and extended strand (17.05%) as generated by PSIPRED (Figure 5).

### 3.5. 3D structure prediction, visualization and physiochemical changes of substitutions

In order to study the biophysical properties of these mutations, Project HOPE server was used to serve this purpose, RaptorX was used to predict a 3D structure model for TP53 protein; while UCSF Chimera was used to visualize the amino acids change (Figure 2). In (Figure 3): (T118L): the amino acid Threonine changes to Leucine at position 118. The mutant residue is bigger than the wild-type residue. The mutant residue is more hydrophobic than the wild-type residue. The residue is located on the surface of the protein, mutation of this residue can disturb interactions with other molecules or other parts of the protein.

**Figure 2:**
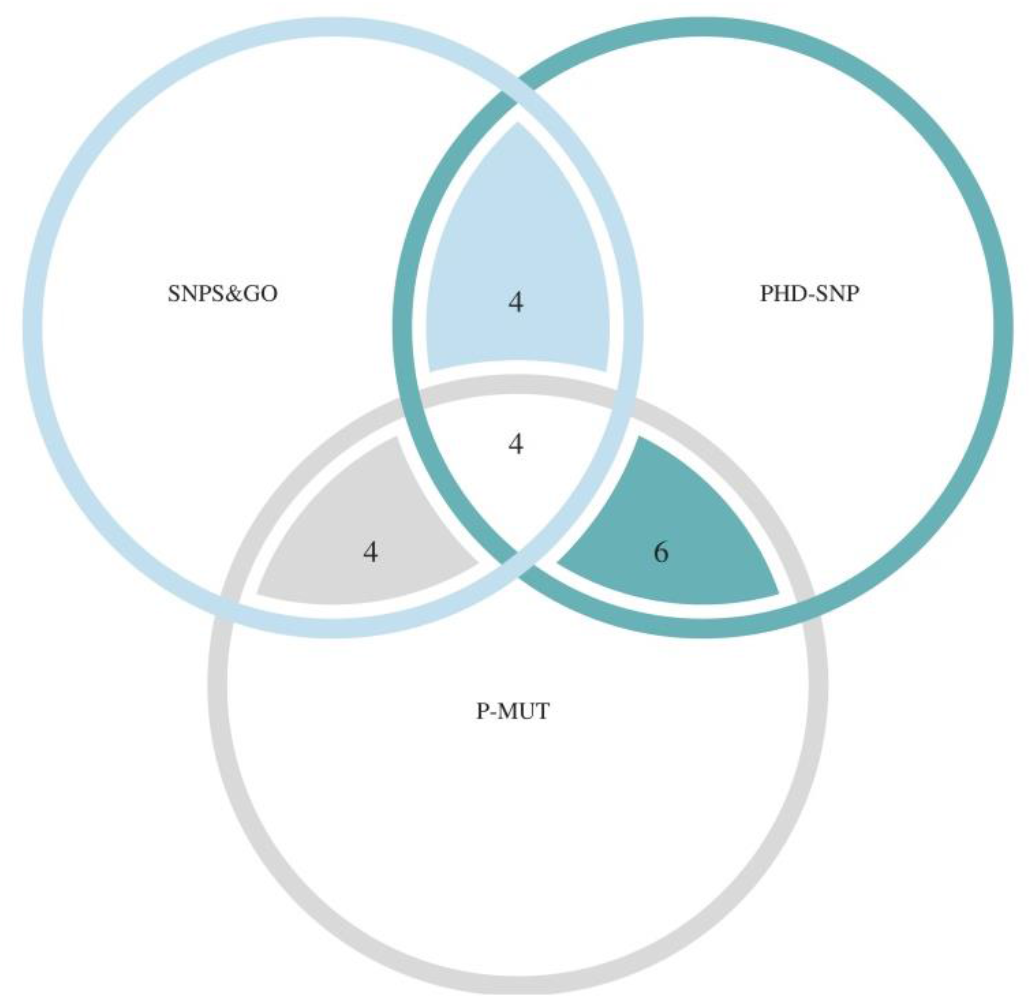
Venn diagram representing the effects of high-risk nsSNPs that shared between SNPs&Go, PHD-SNP, and P-Mut tools.

**Figure 3:**
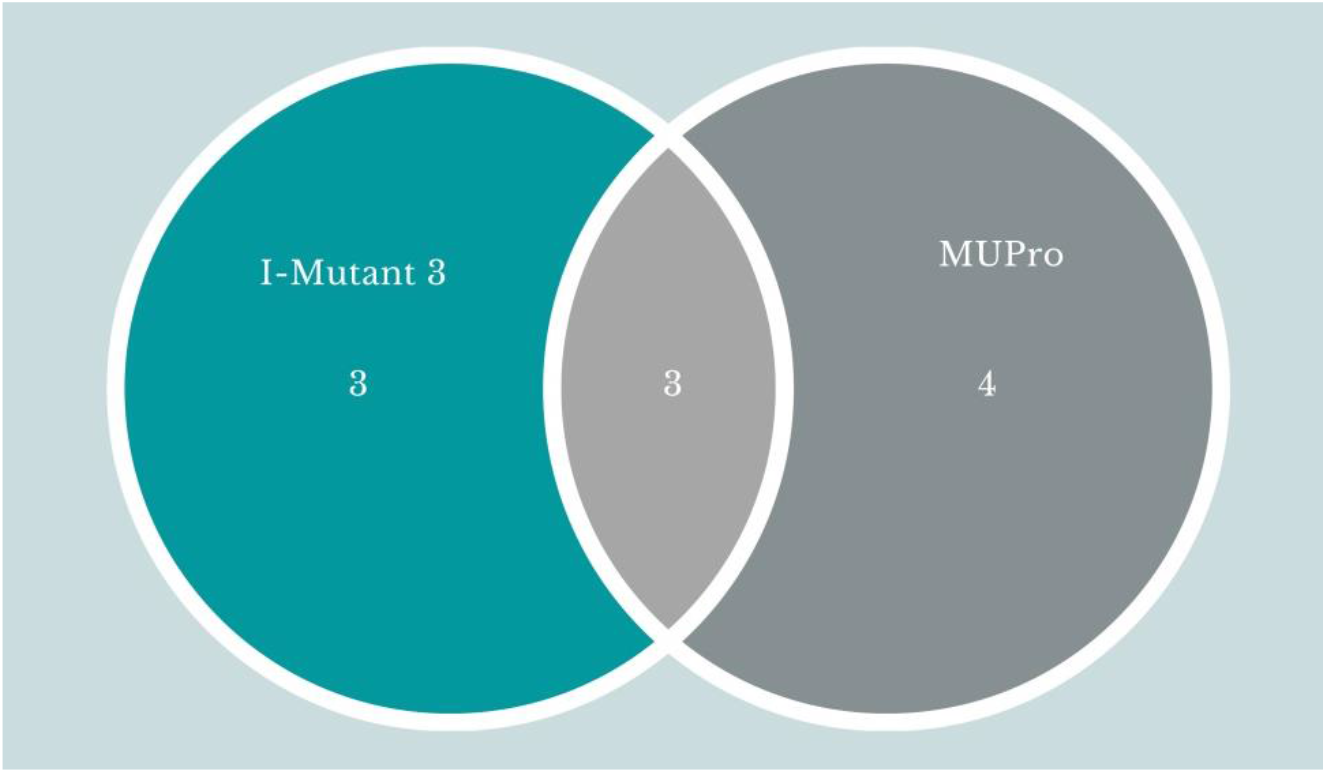
Venn diagram representing the effects of high-risk nsSNPs of PT53 on protein stability shared between I-Mutation 3.0, and MUPro servers.

In (Figure 4): (C242S): the amino acid Cysteine changes to Serine at position 242. The wild-type residue is more hydrophobic than the mutant residue. The mutation will cause loss of hydrophobic interactions in the core of the protein.

**Figure 4:**
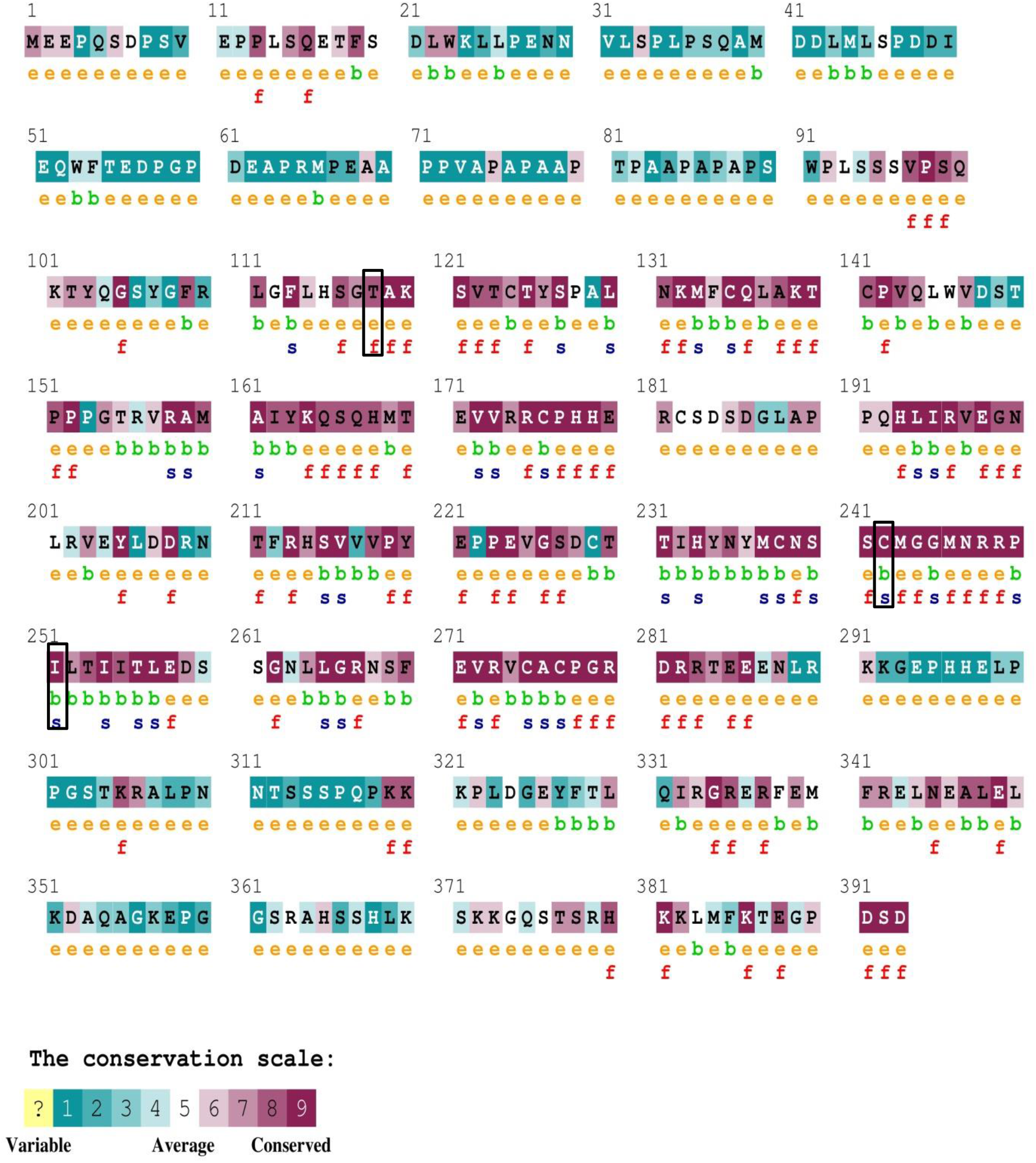
The conserved amino acids across species in TP53 protein were determined using ConSurf. *e*: exposed residues according to the neural-network algorithm are indicated in orange letters. *b*: residues predicted to be buried are demonstrated via green letters. *f*: predicted functional residues (highly conserved and exposed) are indicated with red letters. *s*: predicted structural residues (highly conserved and buried) are demonstrated in blue letters. The black boxes indicate the high-risk nsSNPs.

In (Figure 5): (I251N): the amino acid Isoleucine changes to Asparagine at position 251. The mutant residue is bigger than the wild-type residue. The wild-type residue is more hydrophobic than the mutant residue. The wild-type residue is very conserved.

**Figure 5:**
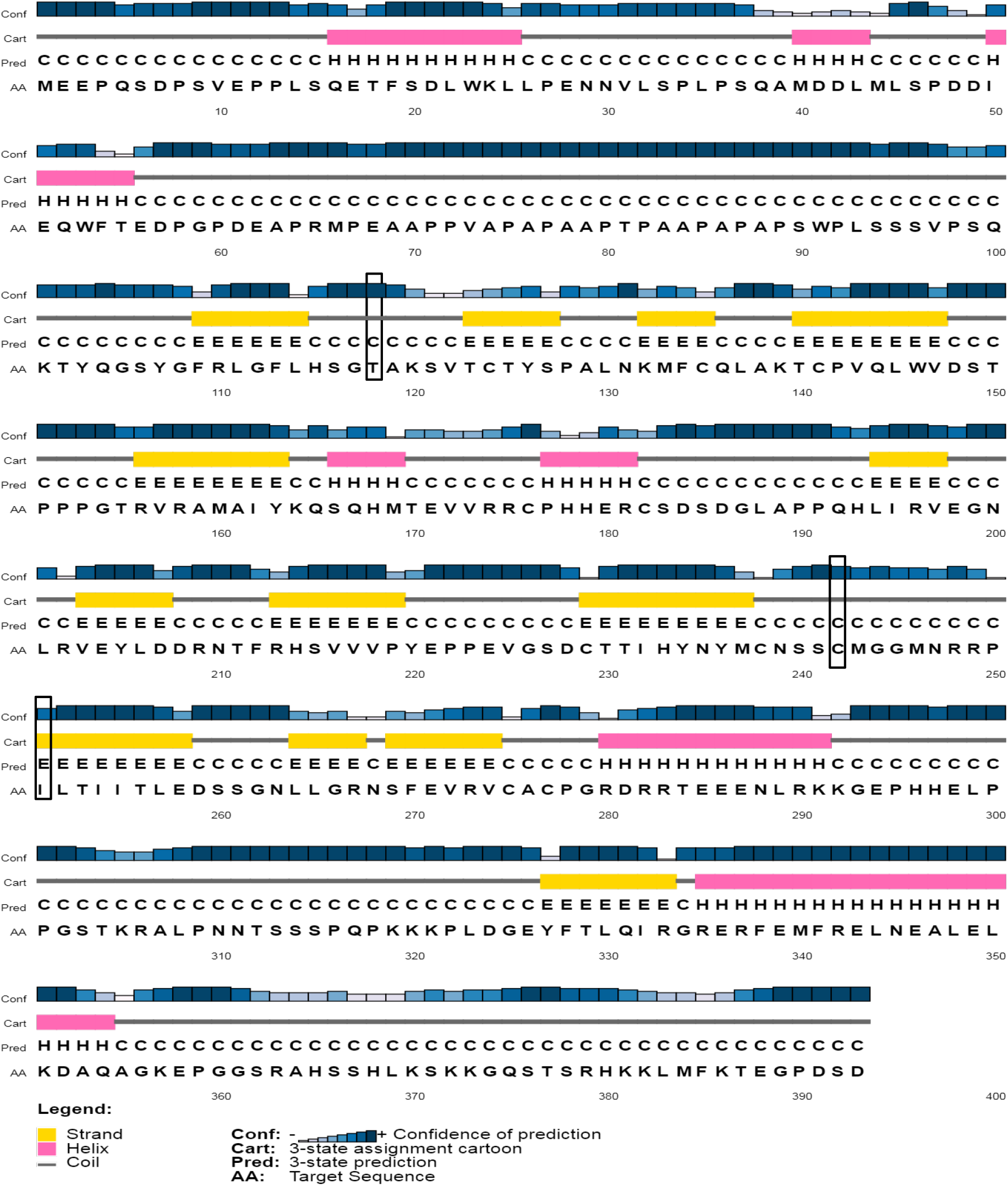
Secondary structure of *PT53* showing beta helix and coil. The results highlight a mix distribution of coil and alpha helix, as generated by PSIPRED server. The black boxes indicate the high-risk nsSNPs.

**Figure 6:**
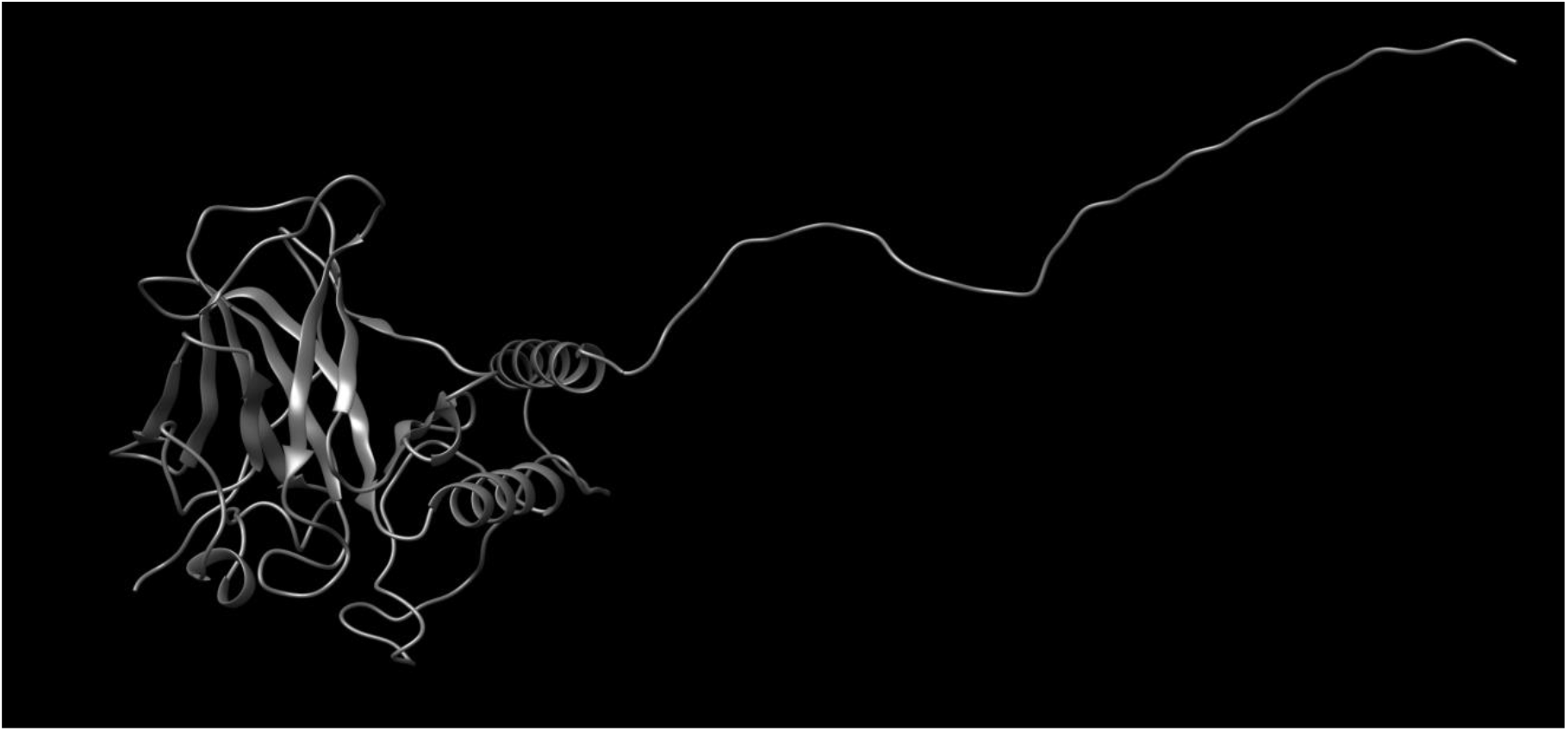
The 3D structure of PT53 protein models by RaptorX.

**Figure 7:**
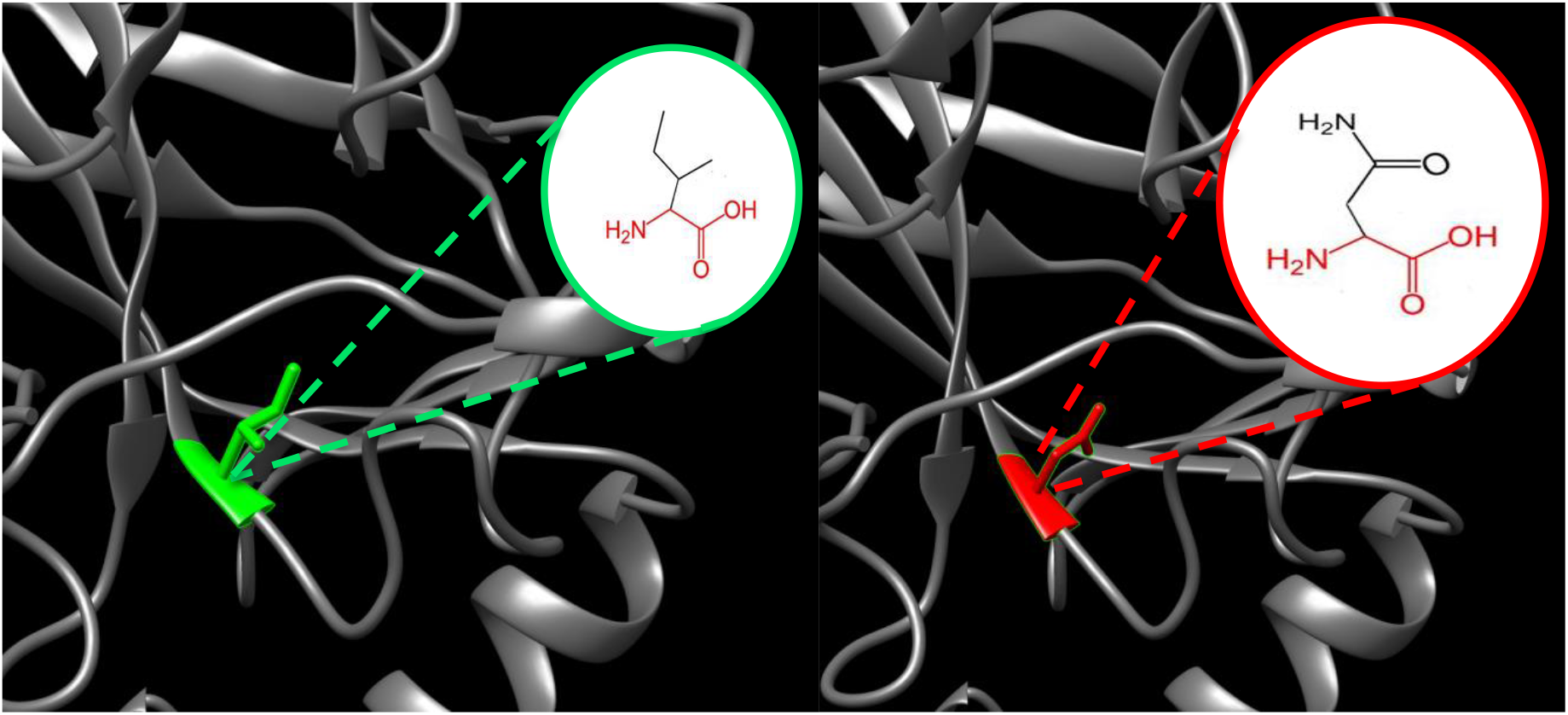
(T118L): The amino acid Threonine changes to Leucine at position 118. The green color indicates the wild amino acid, the red color indicates the mutant one, while the gray color indicates the rest of TP53 protein structure. The 3D observations were done by using UCSF Chimera.

**Figure 8:**
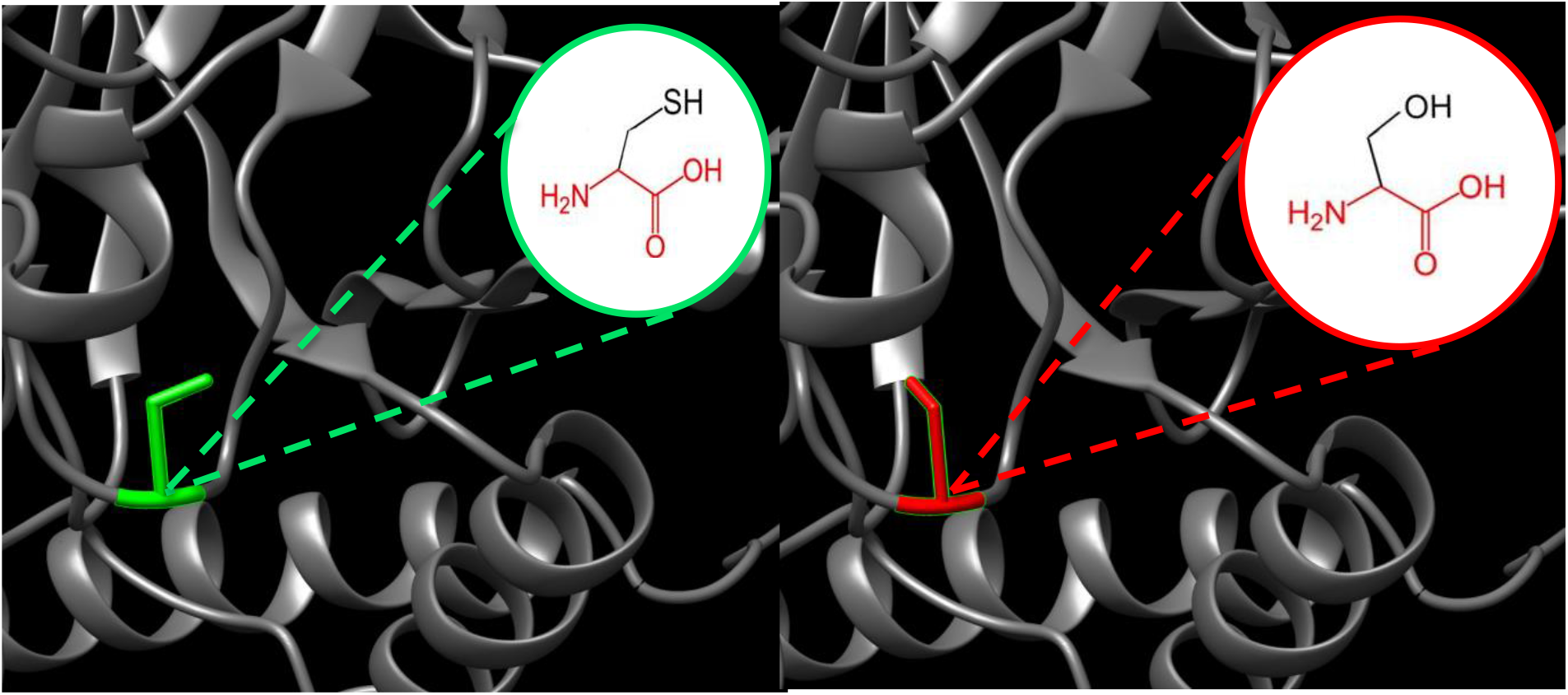
(C242S): the amino acid Cysteine changes to Serine at position 242.

**Figure 9:**
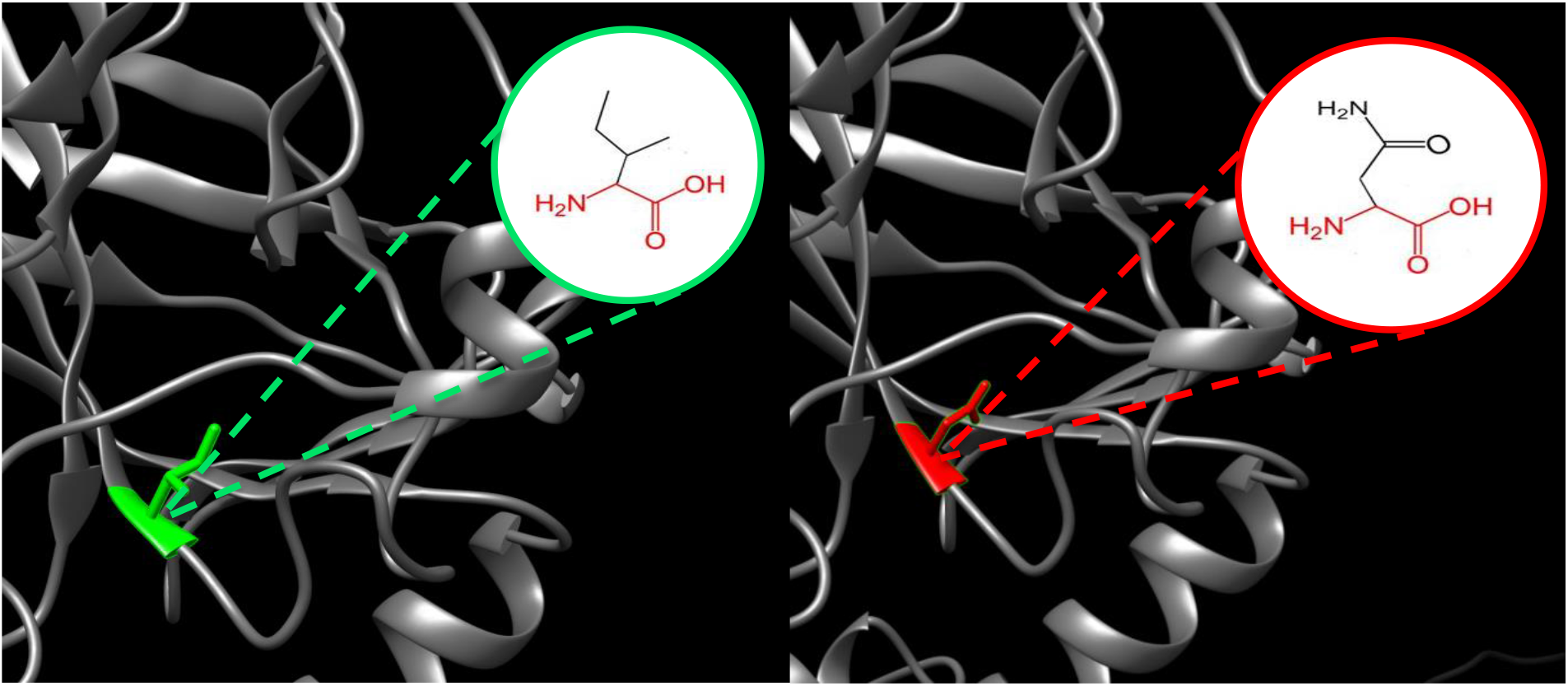
(I251N): the amino acid Isoleucine changes to Asparagine at position 251.

## 4 Discussion

In this study, various sequence-based approaches were performed, because they are more superior to structure-based approaches. Structure-based approaches are restricted to a known 3D structure; on the other hand, sequence-based approaches can be employed in proteins with unknown 3D structures. A combination of multiple predictors revealed better predictions in many recent reports for identifying deleterious nsSNPs in many genes [46-52]. Typically, at least five of *in silico* tools should be considered increasing the consensus on the effect of SNPs [53]; nevertheless, 14 different computational algorithms were engaged to categorize the most pathogenic nsSNPs from *PT53* gene. In the present study a total of three pathogenic SNPs (T118L, C242S, and I251N) were identified after extensive computational analysis.

All three SNPs were also found to be deleterious in common with I-Mutant 2.0 and MUPro servers by destabilizes the protein stability. Furthermore, SNPs that were conserved across the evolutionary perspective are more significant than those that were not conserved, because they are structurally and functionally important for the protein. For this purpose, the ConSurf server was used for evolutionary conservation study. This analysis showed that T118, C242 and I251 were highly conserved in the *PT53* which can directly disturb the protein function and could affect the protein interactions with other proteins.

Our analysis consequently achieves that these SNPs might cause damage to the PT53 protein by disturbing its stability. Nevertheless, the 3D structure of the protein shows a vital role in sympathetic the general effects of nsSNPs on protein function. Thus, 3D protein modeling was achieved by the UCSF Chimera following the improvement of the model to observe the mutational impact on PT53 protein.

In most cases, proteins perform biological functions as temporary or permanent complexes by interacting with other macromolecules. Therefore, mutations in or near some special amino acids that contribute to the functional spatial conformation are at a high risk of causing pathologies. According to the analysis from HOPE, The mutated residues are located in a domain that is important for binding of other molecules. The mutated residues are in contact with a regulatory domain. These mutations could affect this interaction and thereby disturb regulation of the protein.

Also we searched the related published literature to compare our outcomes that had been found by *in silico* method with other approaches, in (I251N) SNP, is present in 0.03% of AACR GENIE (American Association for Cancer Research) cases. This mutation was previously reported to cause lung adenocarcinoma, colon adenocarcinoma, esophageal adenocarcinoma, adenocarcinoma of the gastroesophageal junction, and ampulla of vater carcinoma [54]. The second variant (C242S) was previously reported to cause Acute Myeloid Leukemia [55]; while for the last variant (T118L), we didn’t found any associated clinical studies.

Although it is more reliable to differentiate deleterious SNPs through experiments, it takes much time to execute tests on all variants. Different approaches offer a certain degree of reliability for hazard prediction. By using our computational analysis methods, mainly focuses on 3 high-risk nsSNPs (T118L, C242S, and I251N) from 845. These 3 nsSNPs comes from an systematic analysis summarized in **(figure 10)**. Certainly, there are several limitations of this study; the main limitation is that it focuses on coding region of *PT53* gene using a combination of bioinformatics analysis tools. The second limitation is that the reported causative nsSNPs were limited in number. The third is that there are some similarities of the disease prediction tool between diverse computational analysis tools because most of them were based on changes in conserved residues over time. Nevertheless, *in silico* approach remains a cost-effective technique to achieve a swift analysis concerning the predictable effect of variants.

**Figure 10:**
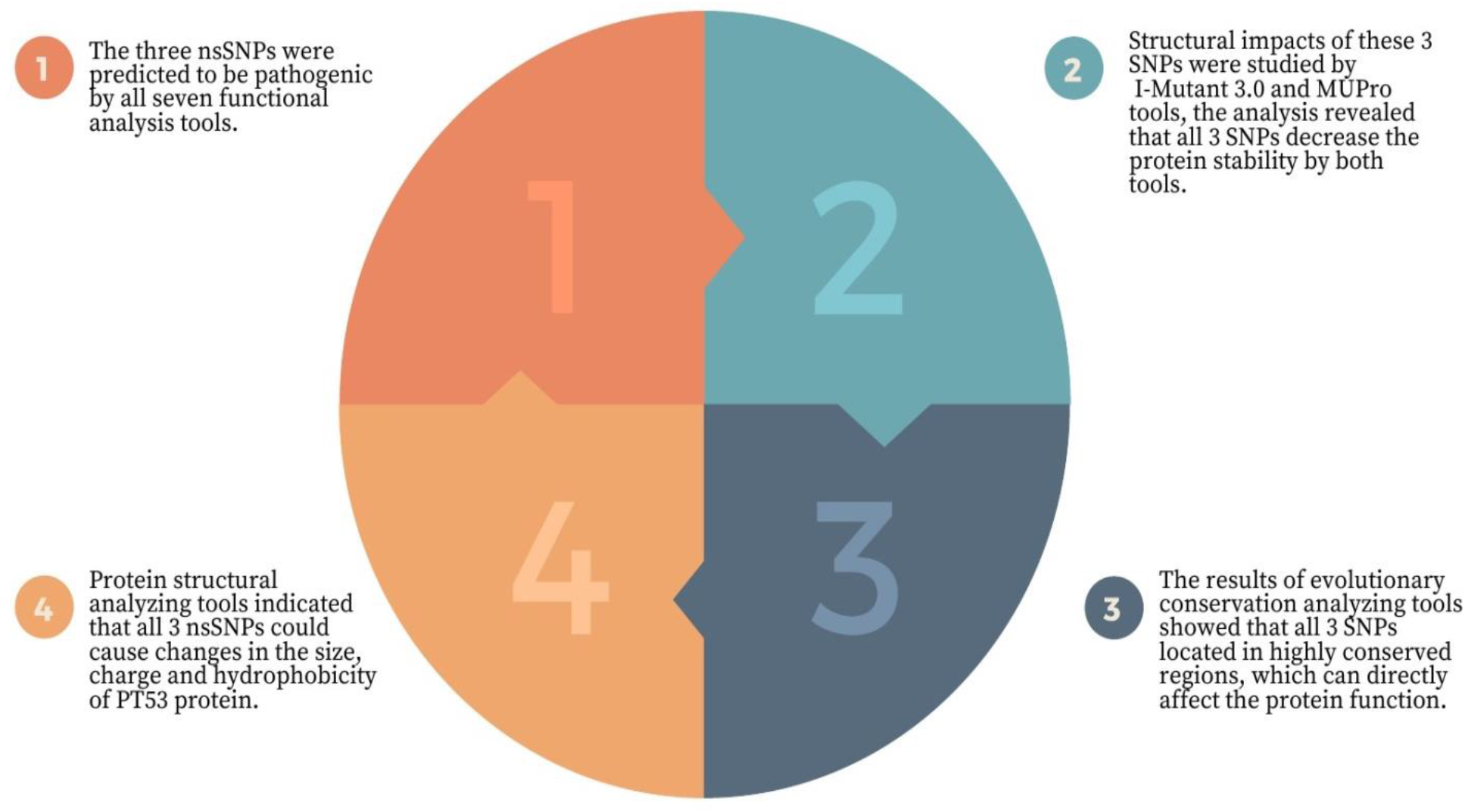
Our strategy to classify the most pathogenic nsSNPs.

## 5 Conclusion

In this study, out of 7 predicted pathogenic nsSNPs, Three high-risk pathogenic nsSNPs of *PT53* gene were identified, the combination of sequence-based and structure-based approaches is highly effective for pointing pathogenic regions. Yet, computational approach can’t replace clinical experiments, and their outcomes should be confirmed by further molecular and clinical studies.

## Data Availability

All data underlying the results are available as part of the article, and no additional source data are required.

## Authors’ Contributions

MIM: conceptualization, formal analysis, investigation, methodology, validation, visualization, and writing (original draft); NSM & MAE: formal analysis, investigation, and validation. AMM: data curation, conceptualization, project administration, supervision, and writing (review and editing). All authors have read and approved the final manuscript.

## Acknowledgments

The authors acknowledge the Deanship of Scientific Research at University of Bahri for the supportive cooperation.

